# Generators and connectivity of the early auditory evoked gamma band response

**DOI:** 10.1101/014514

**Authors:** Nenad Polomac, Gregor Leicht, Guido Nolte, Christina Andreou, Till R. Schneider, Saskia Steinmann, Andreas K. Engel, Christoph Mulert

## Abstract

**Background:** High frequency oscillations in the gamma range are known to be involved in early stages of auditory information processing in terms of synchronization of brain regions, e.g., in cognitive functions. It has been shown using EEG source localisation, as well as simultaneously recorded EEG-fMRI, that the auditory evoked gamma-band response (aeGBR) is modulated by attention. In addition to auditory cortex activity a dorsal anterior cingulate cortex (dACC) generator could be involved. In the present study we investigated aeGBR magnetic fields using magnetoencephalography (MEG). We aimed to localize the aeGBR sources and its connectivity features in relation to mental effort.

**Methods:** We investigated the aeGBR magnetic fields in 13 healthy participants using a 275-channel CTF-MEG system. The experimental paradigms were two auditory choice reaction tasks with different difficulties and demands for mental effort. We performed source localization with eLORETA and calculated the aeGBR lagged phase synchronization between bilateral auditory cortices and frontal midline structures.

**Results:** The eLORETA analysis revealed sources of the aeGBR within bilateral auditory cortices and in frontal midline structures of the brain including the dACC. Compared to the control condition the dACC source activity was found to be significantly stronger during the performance of the cognitively demanding task. Moreover, this task involved a significantly stronger functional connectivity between auditory cortices and dACC.

**Conclusion:** In accordance with previous EEG and EEG-fMRI investigations, our study confirms an aeGBR generator in the dACC by means of MEG and suggests its involvement in the effortful processing of auditory stimuli.

## Introduction

The auditory early evoked gamma-band magnetic fields were discovered more than twenty years ago and localized in the supratemporal auditory cortex (Pantev et al., 1991). The early auditory evoked gamma band response (aeGBR) occurs within 100 ms upon the presentation of an auditory stimulus, and typically consists of oscillations in a frequency range around 40 Hz that can be recorded with electroencephalography (EEG), magnetoencephalography (MEG) and electrocorticography (ECoG). The aeGBR is higher in response to target stimuli than non-target stimuli (Debener et al., 2003), and for louder sounds compared to soft ones (Schadow et al., 2007). Moreover, difficult tasks elicit stronger aeGBR than easy tasks (Mulert et al., 2007). In addition, speech sound aeGBR peaks earlier in the left than in the right hemisphere while completely opposite occurs to non-speech stimuli (Palva et al., 2002). The aeGBR has been increasingly in the focus of interest in recent years, as several EEG (Johannesen et al., 2008; Leicht et al., 2010; Perez et al., 2013; Roach and Mathalon, 2008) and MEG (Hirano et al., 2008) studies have reported reduced power and phase locking of this response in patients suffering from schizophrenia (for an extended review please see (Basar, 2013; Uhlhaas and Singer, 2010; Uhlhaas and Singer, 2013). Given the relevance of GABA-ergic interneurons and the NMDA receptor for gamma oscillations (Fuchs et al., 2007), research into the aeGBR might provide insights for neuropathophysiological mechanisms involved in schizophrenia.

Not only sensory, but also attentional processing has been suggested to contribute to the aeGBR (Leicht et al., 2010; Mulert et al., 2007; Tiitinen et al., 1993). Indeed, a close relationship to selective attention has been described for the 40 Hz aeGBR (Gurtubay et al., 2004; Senkowski et al., 2007; Tiitinen et al., 1997; Tiitinen et al., 1993). Modulation of the aeGBR has been suggested to reflect top-down attentional processing of auditory stimuli (Debener et al., 2003; Schadow et al., 2009). According to the “match-and-utilization” model, the aeGBR is augmented whenever stimulus-related information matches the information loaded into short-term memory (Herrmann et al., 2004). For a review on the association between the aeGBR and attention please see (Fell et al., 2003; Herrmann et al., 2010).

The aeGBR has been invasively recorded in the primary auditory cortex of monkeys (Brosch et al., 2002; Steinschneider et al., 2008) as well as in the primary auditory cortex of neurosurgical patients (Edwards et al., 2005). Furthermore, source localization methods have revealed generators of the aeGBR in the primary auditory cortex (Mulert et al., 2007; Pantev et al., 1991; Schadow et al., 2009) and the corticothalamic network (Ribary et al., 1991) of healthy volunteers. However, an additional generator was suggested to contribute to the aeGBR during demanding, effortful tasks (Ahveninen et al., 2000). This additional generator was indeed later localized in the dorsal anterior cingulate cortex (dACC) using EEG (Mulert et al., 2007) and simultaneous EEG-fMRI (Mulert et al., 2010), which offers high-resolution localization of EEG effects without being limited by the inverse problem (Bénar et al., 2010).

In order to understand the interplay between the dACC and the auditory cortices, it is important to mention evidence from non-human primates suggesting the existence of anatomical connections between the two areas. The dACC receives auditory input from the anterior medial claustrum, while there is a second minor input connection from the auditory association region of the superior temporal area 22 (Mega and Cummings, 1996; Vogt and Pandya, 1987). There are also backprojections extending from the dACC to the association region of the superior temporal area 22 (Pandya et al., 1981). Single trial coupling of the aeGBR and the corresponding BOLD signal revealed the thalamus to be involved in a network including dACC and auditory cortex which is active in correspondence to the aeGBR (Mulert et al., 2010). This might be a correlate of thalamo-cortical interactions (Barth and MacDonald, 1996; Roux et al., 2013) potentially established through cross-frequency coupling with theta oscillations (Buzsaki and Wang, 2012; Jensen and Colgin, 2007; Lisman and Jensen, 2013). However, the cortex is able to generate gamma oscillations independently from subcortical input (Whittington et al., 1995) and corticocortical gamma band interactions have been found in the auditory domain (Bhattacharya et al., 2001; Mulert et al., 2011; Steinmann et al., 2014). Thus and in the light of previous studies suggesting alterations of the aeGBR as an intermediate phenotype of schizophrenia (Leicht et al., 2011), the present study focuses on the aeGBR and its cortical generators.

Several MEG studies have successfully localized various magnetic field responses in the anterior cingulate cortex e.g. (Hirata et al., 2007; Perianez et al., 2004). There have been MEG studies which explored the involvement of dACC in generation of gamma oscillations (Ahveninen et al., 2013; Sedley et al., 2012), however this is the first attempt to explore the potential involvement of the dACC in the generation of the aeGBR.

The aim of the present study was to confirm, using MEG, the presence of the neural generator of the aeGBR in the dACC, and further to investigate how task difficulty influences (1) the aeGBR in the auditory cortex and in the dACC, and (2) the functional connectivity between the two regions.

## Materials and Methods

### Participants

Participants were thirteen right-handed healthy individuals (three females; mean age 25.7 ± 6.5 years, range 18 – 39 years) with no history of neurological or psychiatric disorders. All subjects had audiometric thresholds that were 30 dB HL or better for octave frequencies between 125 to 8000 Hz. The study was approved by the local ethics committee, and informed consent was obtained from all participants prior to inclusion in the study. Participants received a monetary compensation of 15 Euro for the MEG recording session.

### Paradigms

Two versions of an auditory reaction task used in previous studies of our group were administered (Leicht et al., 2010; Mulert et al., 2003; Mulert et al., 2001; Mulert et al., 2005b; Mulert et al., 2007; Mulert et al., 2008), an easy (easy condition, EC) and an attentionally demanding version (difficult condition, DC). In the difficult condition (DC), three tones of different pitch (800 Hz, 1000 Hz and 1200 Hz; 50 repetitions of each) were presented to participants through electrostatic headphones STAX-SR-003 (tube length 1m) in pseudorandomized sequence and interstimulus intervals (ISI: 2.5–7.5 s, mean: 5 s). The duration of each tone was 100 ms, and the sound pressure level was set to 80 dB. Participants were instructed to respond as quickly and as accurately as possible to the low tone by pressing a button with the left index finger, and to the high tone with the right index finger (target tones). The middle tone was not a target for button presses. The easy condition (EC) contained 100 tones with a pitch of 800 Hz (ISI: 2.5–7.5 s, mean: 5 s), to which subjects were requested to respond by left index finger button press. Reaction times (time from stimulus onset to button press) and errors (pressing no button or pressing the wrong button within a timeframe of 150 and 2000 ms after onset of the stimulus) were registered. There was no significant difference between conditions with respect to error rates which were found to be 0.3 % (SD=0.3%) for EC and 4.6 % (SD=5%) for DC. Only trials with correct responses to target tones were considered for further analyses.

### MEG data acquisition

MEG was recorded continuously using a 275-channel (first order axial gradiometers) whole-head system (Omega 2000, CTF Systems Inc.) in a magnetically shielded room. The Presentation^®^ (Version 16.1; Neurobehavioral Systems, USA) software was used for stimulus presentation. MEG data for the two conditions were recorded in two separate consecutive blocks (DC always first). The head position relative to MEG sensors was monitored during each recording block using three fiducial points (nasion, left and right external ear canal). In order to minimize signal incongruence across subjects, special care was taken to place all subjects’ heads inside the MEG dewar in a similar manner.

Maximum head displacement during the recording was 3.2 mm ± 1.9 mm for the DC blocks and 2.8 mm ± 1.5 mm for the EC blocks. There was no significant difference between conditions regarding the amount of head displacement (t[12] = 1.4; p = 0.17).

### Data preprocessing and aeGBR sensor level analysis

Data analysis was conducted in Matlab (MathWorks^®^) using the open source toolbox Fieldtrip (Oostenveld et al., 2011). The MEG signal was low-pass filtered online (cut-off: 300 Hz) and recorded with a sampling rate of 1200 Hz. Offline, the data was segmented into 2-second trials centered around the auditory stimulus onset, filtered low-pass (160 Hz) and high-pass (30 Hz, Butterworth filters order 4; padding with data to 10 sec), filtered for line-noise with band-stop filters for 50, 100, and 150 Hz and demeaned. Trials containing prominent muscle artifacts or sensor jumps were semi-automatically detected and rejected from further analysis. This resulted in 81.8 ± 9.4 average trials per subject in DC and 85.3 ± 7.5 in EC. This is the final number of trials that were used for behavioral and electrophysiological analyses.There was no significant difference between conditions with respect to the number of trials (t[12] = 1.3; p=0.2).

High-pass filtered data are very noisy and only few signal components can be discriminated against this noise. Furthermore and rather independently of the filtering, ICA can be very unstable for a large number of channels. We therefore used PCA (principal component analysis) to robustly reduce data dimensionality and to estimate eigenvalues of the principal components. The PCA was applied to appended trials and eigenvalues of principal components were calculated. Next, an independent component analysis (ICA, infomax extended ICA algorithm with PCA dimension reduction, stop criterion: weight change <10^-7^) was performed on appended trials (Makeig et al., 1996) using only components with an eigenvalue higher than 2.5^-27^, while components with smaller eigenvalues were considered to represent noise. We found this value empirically to result in a stable number of components of around 45 which is large enough to contain most of the signal and small enough to obtain stable ICA components. The average number of calculated independent components per subject was 44 ± 4.5 in DC and 44 ± 4.3 in EC. ICA components representing electrocardiographic artifacts and saccadic spike artifacts were identified and removed considering both topography and time course information. For the correction of saccadic spike artifacts we considered topographies suggested by Carl et al. (Carl et al., 2012).

In order to objectively remove remaining subtle tonic muscle artifacts we developed an algorithm that selected components representing muscle activity based on the power spectrum of each component. It is a well known fact that the power of an EEG/MEG signal approximately corresponds to 1/f (where f stands for frequency), which means that power declines with increasing frequency e.g. (Buzsaki and Draguhn, 2004). This is not the case for muscle activity. Therefore, we correlated the power spectrum of each component derived from a discrete Fourier transform (Hanning taper; 30–120 Hz; 1 Hz steps) with the 1/f function, (for f = 30, 31, 32, ..., 120). Components, for which the Pearson’s correlation index between their power spectrum and the 1/f function was lower than r = 0.7, were considered to be of muscle origin. This way we ensured objectivity in selection of artifactual components, as well as equality of the variance removed in each condition. The final number of rejected independent components was 27 ± 10.8 in DC and 27 ± 9.4 in EC. The remaining components were back-projected to the sensors level and in further text regarded as preprocessed single trials which were used in all analyses.

In order to depict a genuine topographic representation of the aeGBR, preprocessed averaged axial gradiometer data were transformed into planar gradiometer data (Bastiaansen and Knosche, 2000) prior to a time-frequency analysis. Time-frequency analysis was performed using a sliding Hanning-window with 5 ms slide. The length of the time window used for time-frequency analysis was individually calculated for each frequency between 25 and 100 Hz (1 Hz steps; window length = 7/frequency). Subsequently, frequency-wise baseline correction (subtraction) was applied for a pre-stimulus period of 500 ms.

## aeGBR source localization

### Head model

In order to construct head models for source localization, we used individual T1-weighted structural magnetic resonance imaging (MRI) data for 8 participants, while the standard “MNI152” brain template was used for the 5 remaining participants (MRI data not available). Individual MRI data were segmented using the SPM12b software (Friston, 1994), and the grey matter was used for further calculations. Individual head models were subsequently constructed in Fieldtrip using the realistic single shell method (Nolte, 2003).

### Cortical grid and leadfield

Using custom Matlab scripts, an initial cortical grid was created on the cortical surface of the "MNI152" (Mazziotta et al., 2001) brain template as a set of approx. 10000 source points. Of these, we selected 2839 points that were as equally distributed as possible across the brain surface, using an iterative loop: a first source point was randomly selected among the original 10000 source points. For all subsequent source points, we defined the *N*^*th*^+*1* source point as the point with the maximal distance to *N* previously selected source points, which (distance) was defined as the minimum of all *N* distances. The resulting cortical grid was warped for every subject such as to fit the individual MRI. For subjects without individual MRI data, no warping was applied. Every source point represented an equivalent current dipole in the individual source model.

For every subject and condition, the head model, the individual source model and the sensor locations were used to calculate individual leadfield matrices.

### Source localization of aeGBR power

Using the leadfield matrices, the exact low resolution brain electromagnetic tomography (eLORETA) algorithm (as implemented in FieldTrip) was applied in order to produce an eLORETA spatial filter for every leadfield matrix. eLORETA is a discrete, linear, three-dimensional distributed, weighted minimum norm inverse solution, which localizes the power distribution of the MEG signal with exact maxima for single dipoles but with low spatial resolution (Pascual-Marqui, 2007a).

The preprocessed single trials of every subject and each condition were band-pass filtered (35 – 45 Hz; Butterworth filter order 4) and segmented to epochs from 30 to 80 ms (aeGBR segments) and -80 to -30 ms (baseline segments) with respect to the stimulus onset. aeGBR segments were averaged and Fourier transformed (Hanning taper at 40Hz; zero-padding to 2 sec) before calculation of an aeGBR cross-spectrum density (CSD) matrix (according to Equation 1, see below). Our intention was to analyse source power of the aeGBR relative to the pre-stimulus baseline. However, because averaging of baseline activity results in zero values in the denominator of the power ratio, we used the average standard error of baseline single-trial (Fourier transformed; Hanning taper at 40Hz; zero-padding to 2 sec) CSDs as an estimate of the activity of the average background noise. The resulting measure, comparable to the aeGBR, was used to assess relative to baseline gamma response in each condition. The real parts of the baseline CSD matrix and the aeGBR CSD matrix were projected into source space through multiplication with the individual eLORETA spatial filter for every source point. At each source point, power was defined as the maximal eigenvalue of the 3x3 CSD matrix at that source point corresponding to three dipole directions. This value is identical to the power obtained by maximizing power over all possible dipole orientations and assuming fixed orientation during interval of interest over the entire recording block. In order to assess the relative power increase following stimulus presentation compared to the baseline period, the ratio of aeGBR to the estimated baseline activity was calculated for every subject and condition at each source point.

### Regions of interest (ROI) analysis

We used three different ROIs (Fig. 1) corresponding to the left (MNI coordinates: x = -71 to -34; y = -38 to -10; z = 5 to 20) and right (x = 35 to 72; y = -37 to -12; z = 5 to 20) auditory cortices (28 source points each) and the dACC (14 source points, x = -10 to 10; y = 22 to 42; z = 12.3 to 22.3). The source points selected to be included in the left and right auditory cortex ROIs were chosen according to MNI coordinates within Brodmann areas 41 and 42. The coordinates were selected using WFU_pickatlas (Maldjian et al., 2003). The dACC ROI was defined functionally according to the results of a previous study of our group that investigated aeGBR sources using simultaneous EEG and fMRI and the same task as the present study (Mulert et al., 2010): The voxel with the highest gamma-band specific BOLD signal (x = 0; y = 32; z = 22.3; MNI coordinates) in the Mulert et al. (2010) study was used as the center of gravity for the dACC ROI, adding 10 mm in every direction. The resulting dACC ROI included source points from both hemispheres. Additionally, using the WFU_pickatlas (Maldjian et al., 2003), three control ROIs were defined corresponding to the left (MNI coordinates: x = -21 to 0; y = -110 to -82; z = -15 to 10; 23 source points) and right (x = 0 to 21; y = -110 to -82; z = -15 to 10; 19 source points) primary visual cortices and to the posterior cingulate cortex (PCC) (MNI coordinates: x = -16 to 16; y = -52 to -32; z = 10 to 34; 21 source points). The aeGBR power of a ROI was calculated as the mean of the power ratio values of all source points included in the ROI.

**Fig. 1.**
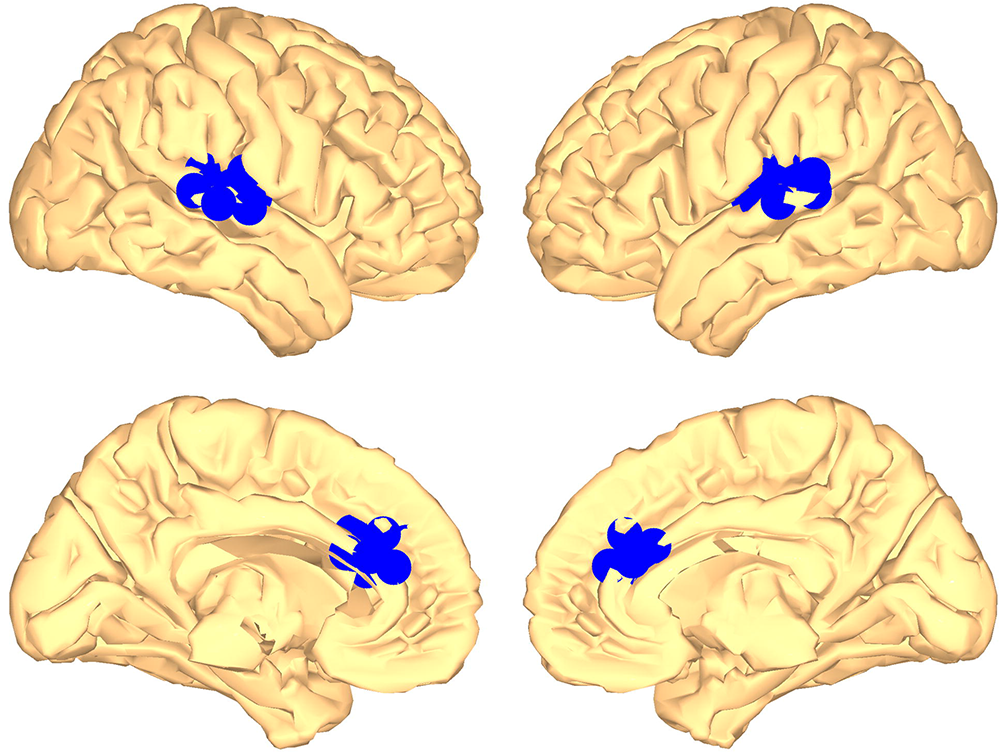
Regions of interest (ROIs) within the left and right auditory cortices (upper row) and the dorsal anterior cingulate cortex (dACC; lower row)

## Estimation of connectivity between ROIs

Functional connectivity between ROIs was estimated using the lagged phase synchronization (LPS) measure (Pascual-Marqui, 2007b). LPS is a measure of dependence between two multivariate time series in the frequency domain based on complex coherency. It has the following properties: a) it vanishes for non-interacting sources regardless of the number of sources and the way they are mapped onto sensors, and b) for any two sources, the actual value is independent of forward mapping. This latter property is the essential difference to using the imaginary part of coherency.

Using the FieldTrip software, preprocessed single trials were band-pass filtered (35 – 45 Hz, Butterworth filter order 4), re-segmented to epochs from 30 to 80 ms after stimulus onset and Fourier transformed (discrete Fourier transformation; Hanning taper at 40Hz with zero-padding to 2 sec). For every MEG sensor, complex frequency domain data were used to build a cross-spectrum density matrix (Equation 1): for two sensors with complex signals *X*(*f*, *i*) and *Y*(*f*, *i*) the cross-spectrum *S*_*XY*_ for a particular frequency *f* at each trial *i* was calculated as:

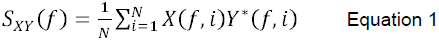

where the asterisk denotes complex conjugate of the complex number, and N refers to the number of trials.

LPS analyses were performed with custom Matlab scripts according to the approach suggested by Pascual-Marqui (Pascual-Marqui, 2007b). In order to map sensor level data onto the source space, we used a new 1-D spatial filter, i.e. we fixed the source orientation which was obtained during eLORETA source localization. This filter was produced by multiplying the eLORETA spatial filter at every source point with the dipole orientation which has the highest power at a particular frequency (in our case 40 Hz). For a detailed account of this filter computation please see (Hipp et al., 2012). Next, the *S*_*XY*_ crossspectrum density matrix was multiplied with that 1-D spatial filter resulting in a source level cross-spectrum density matrix.

The source level cross-spectrum density matrix (s*CSD*, complex valued, size 2839x2839) represents signal interactions among all source points for a particular frequency f. Using this matrix, the complex coherency *Coh* for the frequency f is calculated according to Eq. 2:

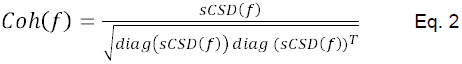

where diag(sCSD) is defined as the diagonal of the sCSD matrix as a column vector, which represents the auto-spectra of the source points, and the square root and the matrix ratio is meant to be elementwise. The transpose of a vector is denoted with the superscript T.

The LPS was calculated according to Eq. 3:

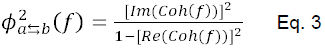

where ◻ represents the LPS, and Im(Coh) and Re(Coh) represent the imaginary and the real part of complex coherency, respectively, between signals at source points a and b for frequency *f*.

The outcome of the Eq. 3 is a 2839x2839 square matrix containing the LPS connectivity values for each pair of source points. The mean LPS values between auditory cortex ROIs and the dACC ROI for each subject and condition were used for the statistical comparison of conditions.

In order to assess the effective connectivity between dACC and auditory cortices the Phase Slope Index (PSI)(Nolte et al., 2008) was calculated for all interactions between voxels included in one of our ROIs and normalized by the respective standard deviation. If the absolute value of the normalized PSI is higher than 2 this connection can be considered as significant. The sign of the normalized PSI reveals the direction of the connection.

## Statistics

All statistical analyses were performed in Matlab (MathWorks^®^). Differences between conditions with respect to head displacement, number of trials, reaction times, relative power increase in ROIs, and LPS values were assessed with dependent sample t-tests. Normal distribution of the respective variables was confirmed with the Shapiro-Wilk parametric test. Correlations between reaction times and aeGBR power in ROIs were assessed using Spearman’s rho.

## Results

There was a significant difference between conditions with respect to reaction times, which were significantly slower in DC (748.2 ± 182.8 ms) compared to EC (350.1 ± 64.2 ms; t[12] = 9.5; p < 0.001).

MEG time-frequency analysis at the sensor level indicated an increase of gamma activity within the aeGBR time window compared to baseline over temporal sensors in both conditions (Fig. 2). Although the scalp topographies of the aeGBR give the impression of a more right dominant aeGBR in EC but not in DC, we did not find any significant side effect. This is in line with previous observations (Ross et al., 2005; Ross et al., 2002).

**Fig. 2.**
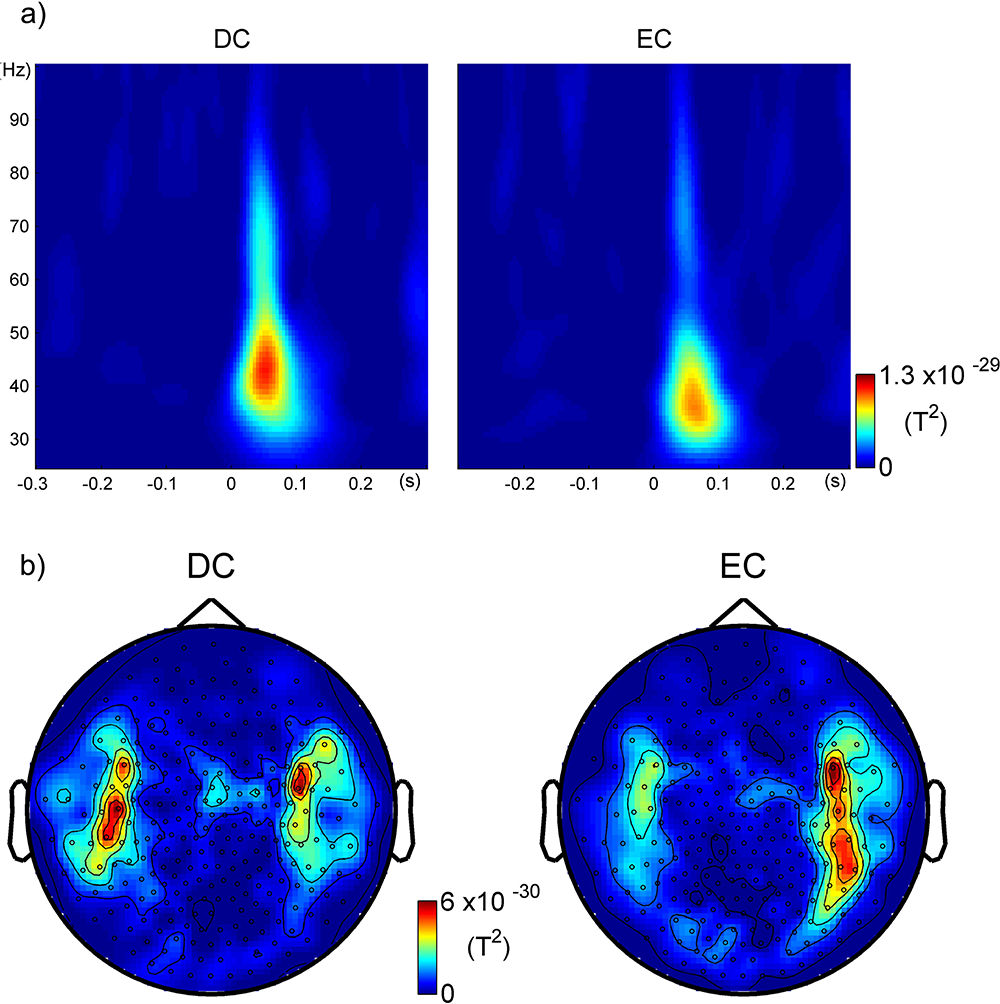
Time frequency plots as a mean over four temporal sensors (MLT13, MLT24, MRT13 and MRT24) and scalp topographies. a) Time-frequency-analysis of the time frame 300 ms before and 300 ms after stimulus presentation averaged over all subjects. Scaling was uniform for both conditions. The early auditory evoked gamma band response (aeGBR) can be seen as an increase of power at approximately 50 ms after stimulus presentation and in the frequency-range around 40 Hz. b) Planar gradients topographies of the aeGBR power for 35 – 45 Hz and 30 – 80 ms after stimulus presentation. DC = difficult condition, EC = easy condition

Source localization of the aeGBR with eLORETA revealed the highest activity within the bilateral superior temporal gyri and in frontal midline structures of the brain including the cingulate gyrus, the medial frontal gyrus and the supplementary motor area in both conditions (Table 1). Inspection of aeGBR activity patterns in the two conditions indicated higher activity in DC than in EC within the dACC (Fig. 3). This impression was confirmed by the ROI analysis, which revealed significantly higher gamma activity in the dACC ROI in DC compared to EC (mean power ratio compared to baseline: 2.6 in DC vs. 1.4 in EC; t[12] = 2.51; p = 0.03; Fig. 4). No significant differences were found between conditions with respect to left and right auditory cortex ROIs.

**Table 1.** Locations of maximum eLORETA activations within different brain regions for the early auditory evoked gamma band response (time frame 30 to 80 ms after stimulus onset), MNI coordinates.

| Condition | Region | x | y | z | Power ratio |
| --- | --- | --- | --- | --- | --- |
| DC | Left supplementary motor area | -3 | -4.1 | 55.1 | 6.22 |
|  | Left cingulate gyrus | -3.7 | -5.5 | 49.1 | 6.21 |
|  | Right supplementary motor area | 3.4 | -5.7 | 56 | 5.91 |
|  | Right cingulate gyrus | 5.5 | -9.1 | 48.1 | 5.82 |
|  | Right superior frontal gyrus | 25.8 | 12.1 | 46 | 5.52 |
|  | Right middle frontal gyrus | 34.6 | 9.9 | 31.6 | 5.47 |
|  | Right postcentral gyrus | 39.7 | -17.7 | 39.3 | 5.13 |
|  | Right superior temporal gyrus | 32.6 | -25 | 14.1 | 5.01 |
|  | Left middle frontal gyrus | -40.6 | 1.6 | 34.5 | 4.81 |
|  | Left superior temporal gyrus | -69 | -35.3 | 17.2 | 4.34 |
| EC | Left supplementary motor area | -1.9 | -11.9 | 53.3 | 5.67 |
|  | Left cingulate gyrus | -1.4 | -11.7 | 40.7 | 5.41 |
|  | Right supplementary motor area | 3.4 | -5.7 | 56 | 5.35 |
|  | Right cingulate gyrus | 5.5 | -9.1 | 48.1 | 5.28 |
|  | Left postcentral gyrus | -42.3 | -10.6 | 34.7 | 5.08 |
|  | Right superior temporal gyrus | 45.6 | -22.3 | 13.6 | 4.91 |
|  | Right precentral gyrus | 41.3 | 7.4 | 27.6 | 4.89 |
|  | Right postcentral gyrus | 44.2 | -9.6 | 34.4 | 4.84 |
|  | Left superior temporal gyrus | -47.2 | -32.7 | 15.7 | 3.28 |
| DC-EC difference | Right superior frontal gyrus | 25.8 | 12.1 | 46 | 2.37 |
|  | Right dorsal anterior cingulate cortex | 5.2 | 21 | 31.6 | 1.86 |
|  | Left superior temporal gyrus | -69.2 | -35.3 | 13.2 | 1.63 |
|  | Left dorsal anterior cingulate cortex | -2.1 | 31.2 | 21.3 | 1.52 |

**Fig. 3.**
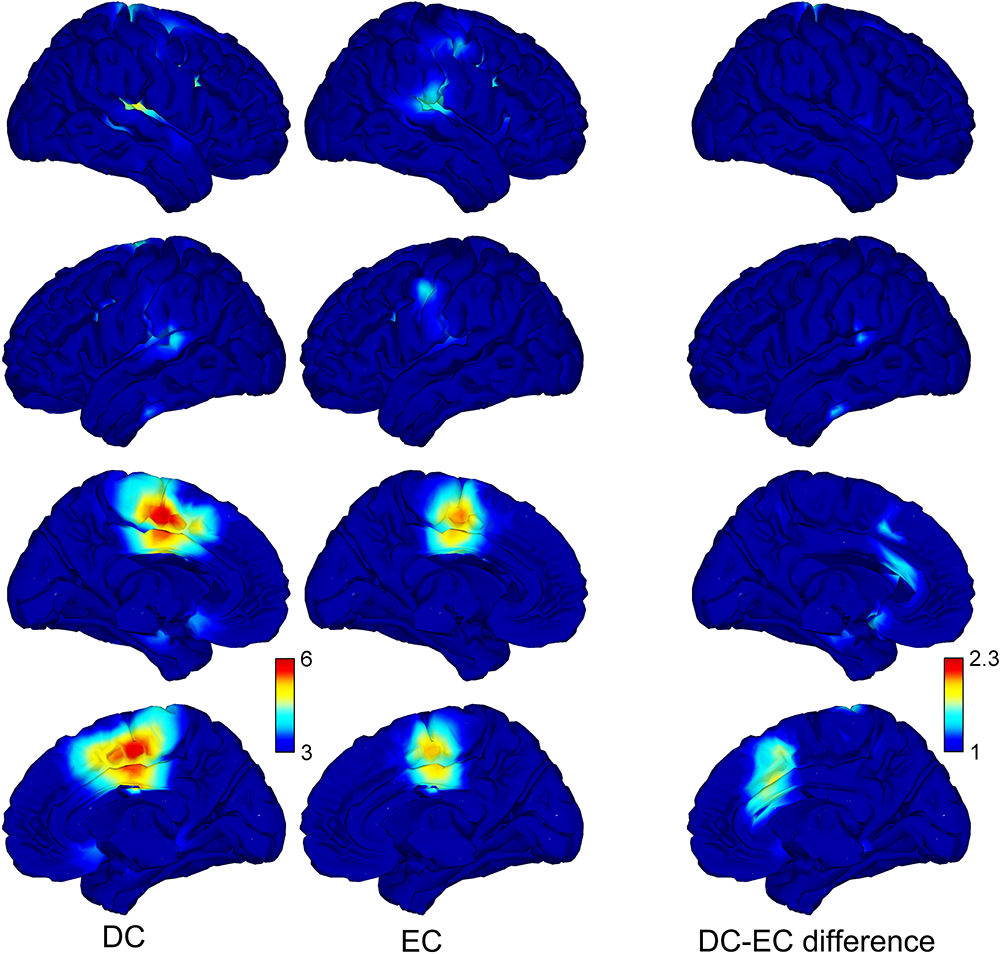
Maximum eLORETA activations for the early auditory evoked gamma band response (aeGBR) within the time frame 30 to 80 ms after stimulus onset presented as power ratio relative to the baseline (mean over all subjects). The aeGBR generators were found within bilateral auditory cortices and midline structures of the brain including the cingulate gyrus, the medial frontal gyrus and the supplementary motor area in both conditions. According to the depicted comparison between conditions (right column) the aeGBR activity of the dACC was stronger during the difficult condition (DC) compared to the easy condition (EC). For plotting purposes we smoothed power ratio values to a 10000 source points grid with a Gaussian spatial filter

**Fig. 4.**
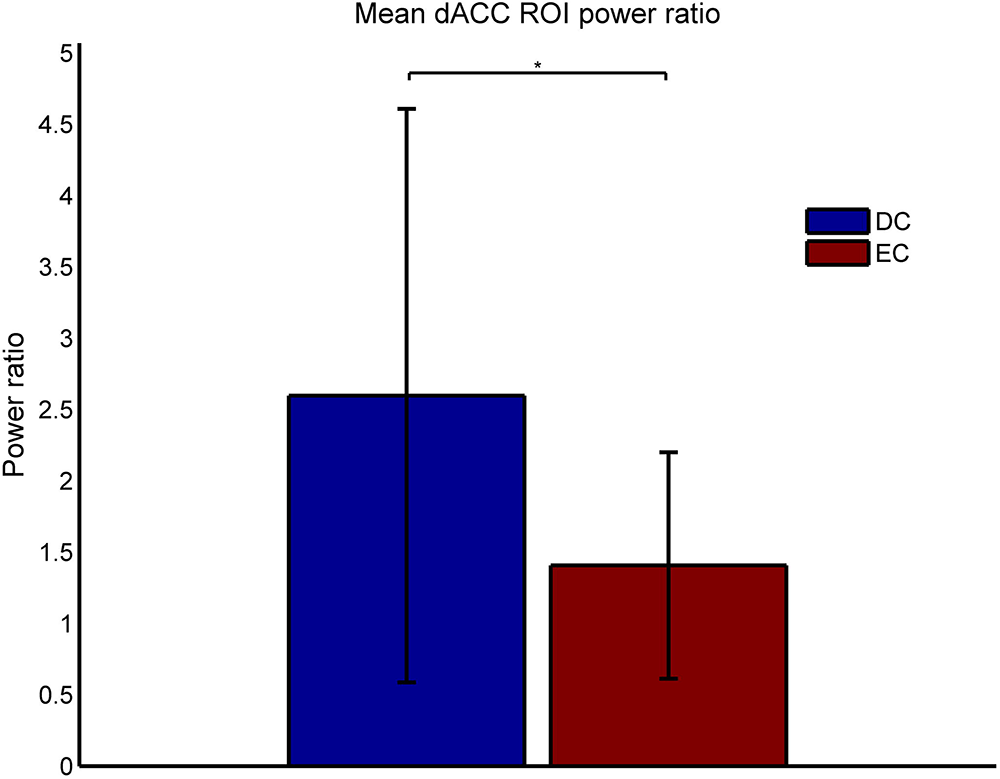
Mean power ratios of early auditory evoked gamma band response (aeGBR) and baseline for the region of interest (ROI) within the dorsal anterior cingulate cortex (dACC). A significantly higher gamma activity within the dACC was found during the difficult condition (DC) compared to the easy condition (EC)

In DC, but not in EC, the activity of the dACC ROI (Spearman’s rho = -0.67, p = 0.01, bootstrap CI= - 0.431 to -0.881) and the mean activity of both auditory cortex ROIs (Spearman’s rho = -0.615, p = 0.03, bootstrap CI = -0.343 to -0.846) inversely correlated with reaction times (Fig. 5). For correlation analysis Fisher z-transformation was applied to ROI activity variables but not to reactions times since this variable was normally distributed.

**Fig. 5.**
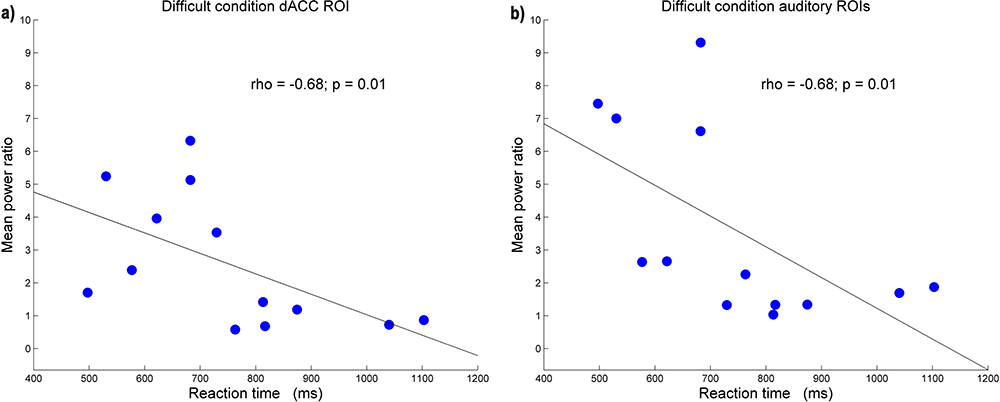
Correlations between reaction times and aeGBR activity within a) the dACC and b) the auditory cortices. The subjects responded faster with increasing mean power ratios between aeGBR and baseline within the dACC region of interest (ROI) as well as within the bilateral auditory cortex ROIs. For correlation analysis Fisher z-transformation was applied to ROI activity variables but not to reactions times since this variable was normally distributed.

Fig. 6 depicts the LPS between the combined bilateral auditory cortices ROIs (seed) and all other source points in the time interval corresponding to the aeGBR. Inspection of the two conditions indicated increased connectivity of the auditory cortex with frontal midline structures of the brain including the dACC in DC compared to EC (Table 2). ROI analysis indicated significantly higher functional connectivity between combined auditory cortices and the dACC in DC (LPS value: 0.06 in DC vs. 0.049 in EC; t[12] = 3; p = 0.01; Fig. 7). There was no significant increase of LPS in DC compared to EC neither between the dACC and the combined control ROIs within the primary visual cortices (t[12] = 1.58; p = 0.14) nor between the auditory cortices ROI and the PCC control ROI (t[12] = -0.1; p = 0.93). The analysis of the PSI failed to give a hint to any directed interaction between the dACC and the auditory cortex.

**Fig. 6.**
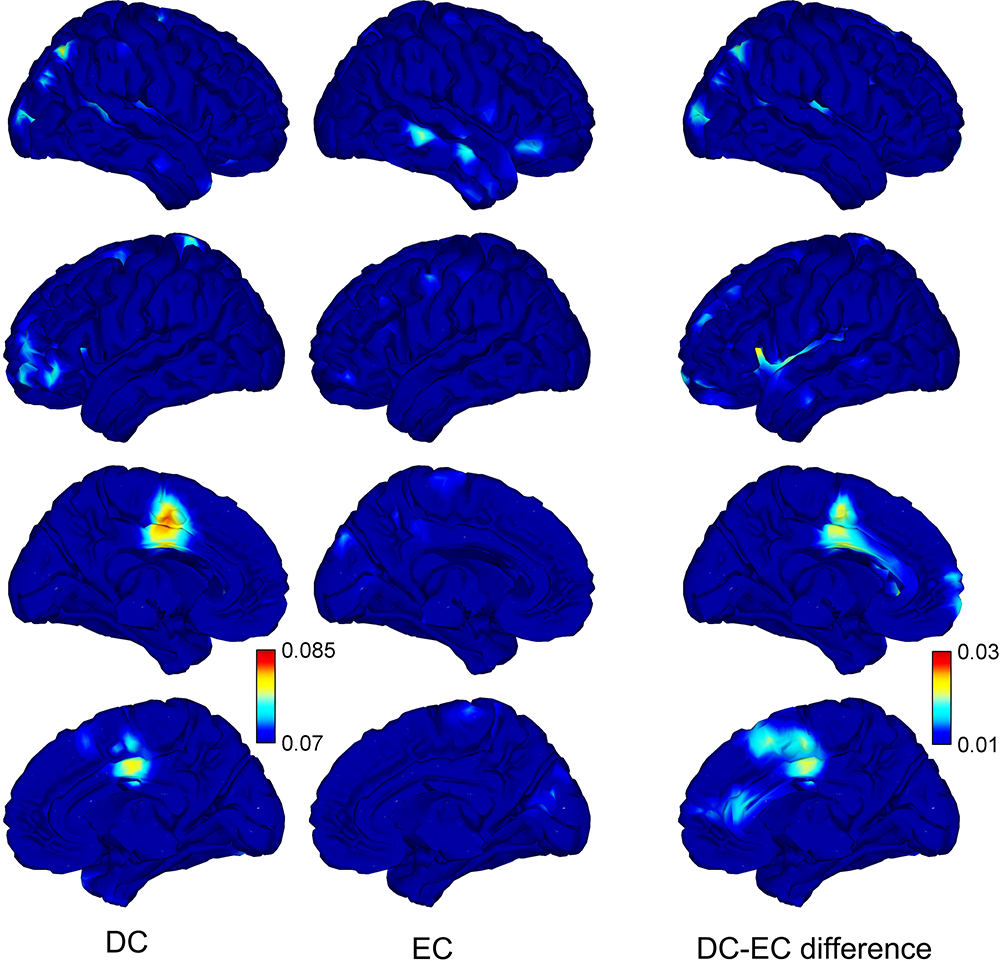
Lagged phase synchronization (LPS) in the gamma-band (35 – 45 Hz) between the bilateral auditory cortices regions of interest as a seed and all other regions of the brain within the time frame 30 to 80 ms after stimulus onset. According to the depicted difference between conditions we observed an increase of connectivity between the auditory cortex and the dACC during the difficult condition (DC) compared to the easy condition (EC). For plotting purposes we used the auditory cortices ROIs as a seed and depicted the LPS values for the connectivity from this seed to the whole brain. LPS values are smoothed to a 10000 source points grid with a Gaussian spatial filter

**Table 2.** Locations of maximum gamma-band lagged phase synchronisation (LPS) between the auditory cortices as a seed and different brain regions within the time window of the early auditory evoked gamma band response (30 to 80 ms after stimulus onset), MNI coordinates.

| Condition | Region | x | y | z | LPS value |
| --- | --- | --- | --- | --- | --- |
| DC | Left supplementary motor area | -8.8 | -2.9 | 48.7 | 0.082 |
|  | Left cingulate gyrus | -3.9 | -4.6 | 43.7 | 0.082 |
|  | Right angular gyrus | 43.5 | -70.4 | 49.3 | 0.079 |
|  | Right cingulate gyrus | 2.5 | -3 | 41.3 | 0.079 |
|  | Left postcentral gyrus | -19.7 | -48.8 | 78.6 | 0.078 |
|  | Right Occipital Lobe (Cuneus) | 26.2 | -88.6 | 7.2 | 0.077 |
|  | Left inferior frontal gyrus left | -48.8 | 42.3 | 4.3 | 0.076 |
|  | Right supplementary motor area | 3.4 | -5.7 | 56 | 0.075 |
| EC | Right middle temporal gyrus | 71.3 | -35.9 | -2.2 | 0.077 |
|  | Right middle temporal gyrus | 67.9 | -10.2 | -13.9 | 0.075 |
|  | Right inferior frontal gyrus | 50.6 | 35.1 | -13.4 | 0.074 |
| DC-EC difference | Left insula | -33.7 | 5.6 | 13.3 | 0.026 |
|  | Left supplementary motor area | -3 | -4.1 | 55.2 | 0.022 |
|  | Left cingulate gyrus | -4 | -4.6 | 43.7 | 0.021 |
|  | Right cingulate gyrus | 2.5 | -3 | 41.3 | 0.020 |
|  | Right supplementary motor area | 3.4 | -5.7 | 56.1 | 0.019 |
|  | Right angular gyrus | 43.5 | -70.4 | 49.4 | 0.019 |
|  | Right occipital pole | 32.2 | -95.2 | 11.4 | 0.018 |
|  | Right anterior cingulate cortex | 60 | 32.5 | 11.6 | 0.017 |

**Fig. 7.**
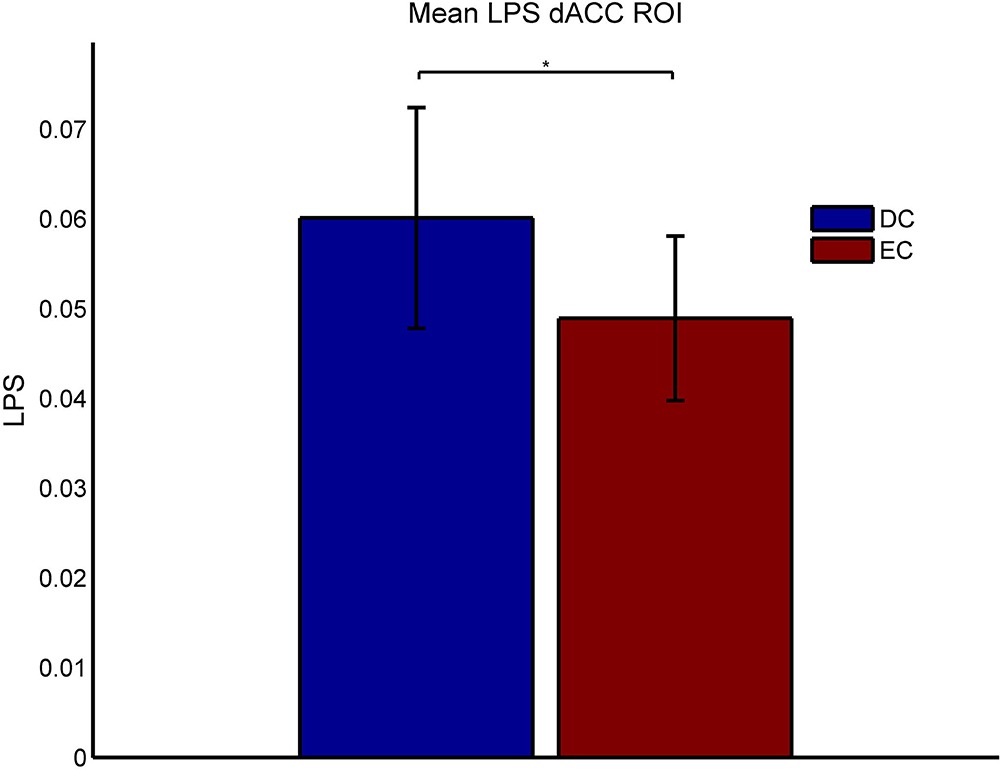
Lagged phase synchronization (LPS) values representing the functional connectivity between dACC and bilateral auditory cortices (mean over all subjects). The bilateral auditory cortices regions of interest (ROIs) were used as a seed. During occurrence of the early auditory evoked gamma band response (30 – 80 ms after stimulus onset) the difficult condition (DC) shows significantly stronger functional connectivity between auditory cortices and dACC compared to the easy condition (EC)

## Discussion

The present study used MEG to investigate the aeGBR in healthy subjects. The two versions of the paradigm differed with respect to their demands on the cognitive system (Mulert et al., 2007), as in DC subjects had to distinguish among three different tones. Consistent with our hypotheses, we found MEG neural generators in bilateral auditory cortices and in frontal midline structures of the brain. The auditory sources of the aeGBR show some differences between conditions with respect to their location within the auditory cortex. Under restriction of the spatial resolution derived from the MEG-based source localisation one could speculate that rather primary auditory cortex regions were found during EC and rather secondary auditory cortex regions (BA22) during DC. The frontal midline generator was present during both conditions (see Fig. 3, first and second column), but it showed substantially stronger activity in the difficult condition. According to the comparison between conditions (Figure 3, third column), the region that contributed most to the frontal midline generator was the dACC, while ROI analysis confirmed significantly greater aeGBR power increase in the dACC in difficult compared to the easy condition (Fig. 4). Functional connectivity analyses also showed significantly stronger functional connectivity between the auditory cortices ROIs and the dACC ROI in the difficult compared to the easy condition (Fig. 6). This was specifically the case for the connection between auditory cortex and dACC as no increase of LPS was present between dACC as well as auditory cortex and control ROIs. Analysis of behavioral results (Fig. 5a) suggested an inverse correlation between aeGBR power in the dACC and reaction times in the difficult, but not in the easy condition. The same inverse correlation pattern was observed for aeGBR power in auditory cortices and reaction times in the difficult condition (Fig. 5b). Our results are in line with findings from previous EEG (Mulert et al., 2007) and coupled EEG-fMRI (Mulert et al., 2010) studies, and provide MEG evidence in support of the hypothesis that the dACC is implicated in effortful processing of auditory stimuli. This appears to be a specific effect of the dACC, as we found no difference in aeGBR between conditions for the auditory cortex ROIs.

Notably, the aeGBR shows different spectral peaks with respect to our different conditions (Fig. 2a) This might give a hint to an influence of increasing cognitive demands on the peak frequency of evoked gamma oscillations which has to be explicitly addressed in future studies.

The dACC is a multifunctional structure, involved in a variety of cognitive tasks. Being the subject of a large body of neuroscientific research, this region has been associated with a variety of functions, e.g. acceleration of reaction times (Bush et al., 1998), conflict monitoring (Botvinick et al., 1999), response selection (Turken and Swick, 1999), reward-based decision-making (Bush et al., 2002), error detection (Fiehler et al., 2004), mediation of behavioral adaptation based on the assessment of outcome of own actions (Sheth et al., 2012), selection and maintenance of behaviors (Holroyd and Yeung, 2012), and action initiation (Srinivasan et al., 2013). For a comprehensive review of the functional neuroanatomy of cingulate cortex please see (Vogt, 2009). Given the variability of the suggested functions of the dACC, considerable efforts have been made to unify these functions into one single model that could reconcile results across different research fields and paradigms. Functional neuroimaging studies have typically interpreted the function of the dACC mainly through its involvement in cognitive processes and reinforcement learning, while neuropsychological studies in patients with cerebral lesions have suggested a role of the dACC in motivating and "energizing" behavior (Holroyd and Yeung, 2012).

It is of special interest to shortly discuss how recent models of dACC function explain its involvement in a mental effort demanding task in the present study. Holroyd & Yeung (2012) suggested "the actor-critic model” that draws upon hierarchical reinforcement learning theory. According to these authors, the dACC is part of a larger behavioral network that plays a role in selection and maintenance of a given action (rather than conflict monitoring), as well as in sustaining effortful behavior (Holroyd and Yeung, 2012). Another relevant model proposed by Shenhav et al. (2013) postulates an involvement of the dACC in cognitive control through (a) coding the “expected value” (payoff) of controlled processes, (b) computing the amount of control that needs to be invested in order to achieve this payoff, and (c) estimating the cost in terms of cognitive effort required (Shenhav et al., 2013). According to this model, the role of the dACC is to maximize the expected value of control, taking into account the above three variables. In line with these models of ACC function previous EEG and simultaneous EEG-fMRI studies using the same or slightly modified tasks as compared to the present study have reported significant associations between dACC activity and conscious mental effort (Mulert et al., 2005a; Mulert et al., 2008) as well as passive mental effort demands (Mulert et al., 2010; Mulert et al., 2007). The latter investigated the dACC as a generator of the aeGBR. Moreover, subjects performing the task used here showed increased N100 amplitude and dACC N100 source activity in relation with faster reaction times (Mulert et al., 2003; Mulert et al., 2005a). A similar association was noted in the present study between dACC aeGBR activity and improved performance. Thus, the present study confirms and expands previous EEG findings using an MEG-based approach.

Accordingly, the location of the dACC ROI used in the present study was chosen based on our previous work investigating the aeGBR with simultaneous measurement of EEG and fMRI (Mulert et al., 2010). This location is well in line with the location of the maximum difference between conditions within the ACC obtained from the eLORETA aeGBR power analysis (see Fig. 3), although a more posterior location of ACC activity is visible in the separate analysis of conditions. Given the resolution of the MEG the analysis of dACC and auditory cortex ROIs might have been confounded by activity of surrounding brain regions. However, both EEG-based source localization (Leicht et al., 2010; Mulert et al., 2007) and single trial EEG-fMRI coupling (Mulert et al., 2010) suggested generators of the aeGBR within the dACC and the bilateral auditory cortices.

As a cautionary note, it should be mentioned that our findings do not constitute direct proof for a top-down involvement of the dACC. They do nevertheless suggest an early synchronization of sensory and monitoring regions during the effortful processing of auditory stimuli. According to the “match and utilization model”, stimuli that match previous memory content (in this case, target stimuli) are more salient than non-target stimuli (Herrmann et al., 2004). Selective attention towards a relevant feature of stimulus (in this case, pitch frequency) leads to generation of top-down signals that drive the synchronization of sub-threshold oscillations of feature-selective neural assemblies. Therefore, the augmentation of the aeGBR, generated in the auditory cortices and the dACC, might represent the means through which selective attention exerts its top-down influence on the primary auditory cortex. Although the aeGBR, as an evoked (time-locked) response, corresponds to an average that disregards single trial phase information needed for connectivity analyses, one might argue that all regions that are active during the time-locked “event” are conceivably connected with each other. This notion was confirmed in the present study by looking into functional connectivity patterns of the 40 Hz total gamma signal at the time interval where the aeGBR was the most prominent, i.e. 30 – 80 ms after the stimulus. However, it is important to mention that lagged phase synchronization conveys information regarding functional connectivity, but not its direction. Unfortunately, our attempt of estimating the direction of interaction between dACC and auditory cortex using the Phase Slop Index failed to reveal significant results which might be due to an undersized sample. A limitation of our study is the fact that an increase in gamma power in DC compared to EC could result in an increase of the LPS measure due to an enhanced signal to noise ratio. However, in our study, LPS values were not correlated with power ratio values in each of the conditions. Based on this analysis, we can exclude linear relationships between the two variables whereas there is no way to exclude non-linear relationships.

The present findings are in line with the notion of gamma-band long-range connectivity (Buzsaki and Wang, 2012; Singer, 1999; Uhlhaas and Singer, 2013). The long-range gamma band synchronization has been shown to be mediated by theta oscillations (Jensen and Colgin, 2007; Lisman and Jensen, 2013) and oscillations from other frequency ranges such as delta, beta and alpha oscillations (Canolty and Knight, 2010; Fontolan et al., 2014; Roux et al., 2013). However, so far, there are no studies explicitly investigating the relation between the aeGBR and other frequency ranges.

A limitation of our study is that we did not detect signals from deep sources such as the thalamus, which has been suggested to be involved in the generation of the aeGBR in previous studies (Mulert et al., 2010). However, the question whether MEG has the capacity to detect signals from such deep sources is still under debate (Roux et al., 2013; van Wijk and Fitzgerald, 2014). One possibility is that the functional coupling between the auditory cortex and the dACC observed in the present study is partly mediated by the thalamus. This interpretation would be consistent with the suggestion that functional coupling in the 40-Hz range across long distances requires strong anatomical connections, e.g. large diameter myelinated fibers with satisfactory conduction speed (Aboitiz et al., 2003).

In conclusion, our study confirmed an MEG generator of the aeGBR in the dACC. Increased activity of this 40Hz generator was associated with faster reaction times, dependent on the demands placed by the sensory task on the cognitive system. In the same time window and frequency, functional connectivity between auditory cortices and the dACC was increased with mental effort.

The authors declare no conflict of interest. This work has been supported by DFG, SFB 936 "Multi-Site Communication in the Brain", project (SFB936/C6/A3/Z1).

